# Mitochondria are physiologically maintained at close to 50 °C

**DOI:** 10.1101/133223

**Authors:** Dominique Chrétien, Paule Bénit, Hyung-Ho Ha, Susanne Keipert, Riyad El-Khoury, Young-Tae Chang, Martin Jastroch, Howard T Jacobs, Pierre Rustin, Malgorzata Rak

**Affiliations:** INSERM UMR1141, Hôpital Robert Debré, 48, Boulevard Sérurier, 75019, Paris, France; Université Paris 7, Faculté de Médecine Denis Diderot, Paris, France; College of Pharmacy, Suncheon National University, Suncheon, 540-742 Republic of Korea; Institute for Diabetes and Obesity, Helmholtz Centre Munich, German Research Center for Environmental Health, 85764 Neuherberg, Germany; Neuromuscular Diagnostic Laboratory, Department of Pathology & Laboratory Medicine, American University of Beirut Medical Center, Beirut, Lebanon; Department of Chemistry, POSTECH, Pohang, Gyeongbuk, 37673 Republic of Korea; BioMediTech and Tampere University Hospital, FI-33014 University of Tampere, Finland; Institute of Biotechnology, FI-00014 University of Helsinki, Finland

## Abstract

In endothermic species, heat released as a product of metabolism ensures stable internal temperature throughout the organism, despite varying environmental conditions. Mitochondria are major actors in this thermogenic process. Part of the energy released by the oxidation of respiratory substrates drives ATP synthesis and metabolite transport, while a noticeable proportion is released as heat. Using a temperature-sensitive fluorescent probe targeted to mitochondria, we measured mitochondrial temperature *in situ* under different physiological conditions. At a constant external temperature of 38 °C, mitochondria were more than 10 °C warmer when the respiratory chain was fully functional, both in HEK293cells and primary skin fibroblasts. This differential was abolished in cells lacking mitochondrial DNA or by respiratory inhibitors, but preserved or enhanced by expressing thermogenic enzymes such as the alternative oxidase or the uncoupling protein 1. The activity of various RC enzymes was maximal at, or slightly above, 50 °C. Our study prompts a re-examination of the literature on mitochondria, taking account of the inferred high temperature.

## Introduction

As the main bioenergetically active organelles of non-photosynthetic eukaryotes, mitochondria convert part of the free energy released by the oxidation of nutrient molecules into ATP and other useful forms of energy needed by cells. However, this energy conversion process is far from being 100% efficient and significant fraction of the released energy is dissipated as heat. This raises the hitherto unexplored question of the effect of this heat production on the temperature of mitochondria and other cellular components.

To address this issue, we made use of the recently developed, temperature-sensitive fluorescent probe (Fig. S1A), MitoThermo Yellow (MTY) [1]. Because molecule fluorescence is known to be quite sensitive to a number of factors and as MTY is derived from the membrane potential-sensitive dye rhodamine, in this study we investigated whether the observed changes in MTY fluorescence we observed in HEK293 cells (human embryonic kidney cells 293) could be influenced by altered membrane potential, or by associated parameters, such as pH, ionic gradients or altered mitochondrial morphology. As a major conclusion of this study, we found that the rise in mitochondrial temperature due to full activation of respiration is as high as ∼10 °C (n=10, range 7-12 °C; compared to 38°C temperature of cell suspension medium). We also showed that respiratory chain activities measured in intact mitochondria are up to 300% increased when assayed at the mitochondrial temperature measurable in intact cells.

## Results

We first confirmed targeting to mitochondria in both HEK293 cells and primary skin fibroblasts, based on co-localization with the well-characterized dye MitoTracker Green (MTG; Fig. 1A). It was previously shown that the initial mitochondrial capture of MTY was dependent on the maintenance of a minimal membrane potential [1]. The exact sub-mitochondrial location of the probe is yet to be established, although it has been postulated to reside at the matrix side of the inner membrane [1]. MTY fluorescence from mitochondria was retained over 45 min, regardless of the presence of respiratory chain inhibitors, whilst full depolarization with an uncoupler led to MTY leakage from mitochondria after only 2 min (Fig. S3). Fluorescence remained stable over 2 hours in HEK293 cells, although the degree of mitochondrial MTY retention varied between cell lines, with probe aggregation observed in the cytosol in some specific lines (Fig. S3A). In HEK293 cells, which were selected for further study, we observed no toxicity of MTY (100 nM in culture medium) over 2 days (Fig. S5).

**Figure 1.**
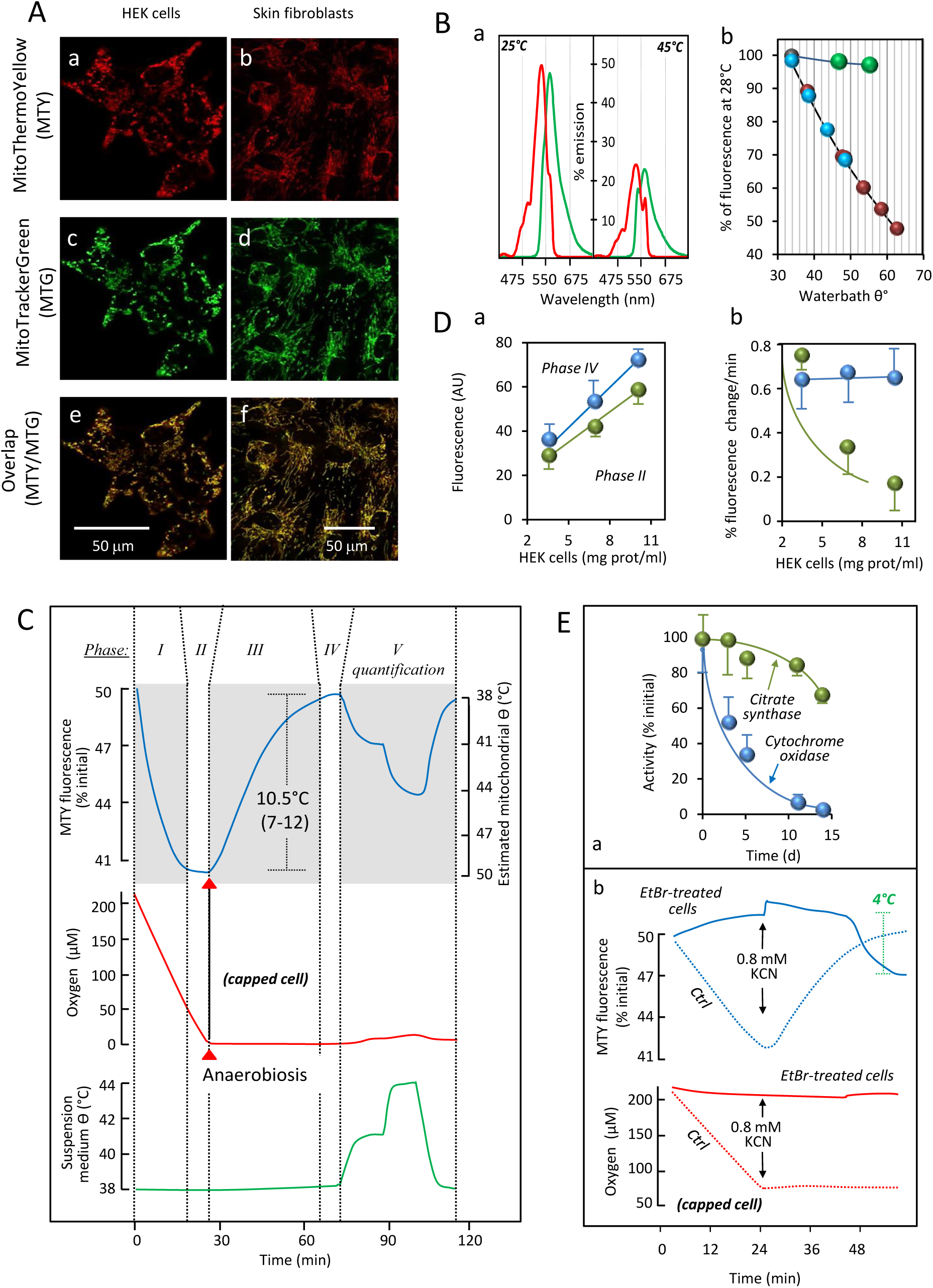
Determination of mitochondrial temperature in intact human cells. A: The temperature-sensitive probe MitoThermo Yellow (MTY: a, b) co-localizes with MitoTracker Green (MTG: c, d, merge: e, f) in HEK293 cells and in primary skin fibroblasts, as indicated. B: a, Fluorescence excitation (red) and emission (green) spectra of MTY (1 mM) in 2 ml PBS at 25 and 45 °C; b, Response of MTY (blue and red) and rhodamine (green, also 1 mM) fluorescence in 2 ml PBS to temperature (34 to 64 °C). Note that the pseudo-linear decrease of MTY fluorescence corresponding to increasing temperature (blue) is essentially reversed upon cooling (red) of the solution to the initial temperature. C, The definition of the various phases of fluorescence in MTY-preloaded HEK293 cell adopted in this study. Note that the initial value is given systematically as 50%, as set automatically by the spectrofluorometer, allowing either increases or decreases to be recorded. Phase I: cell respiration (red trace) after cells are exposed to aerobic conditions in PBS, resulting in decreased MTY fluorescence (blue trace) as mitochondria heat up; phase II: cell respiration under aerobic conditions, where a steady-state of MTY fluorescence has been reached (maximal warming of mitochondria). Cells were initially maintained for 10 min at 38 °C under anaerobic conditions, before being added to the cuvette; phase III: cell respiration that has arrested due to oxygen exhaustion -MTY fluorescence progressively increases to the starting value, as mitochondria cool down; phase IV: stalled respiration due to anaerobiosis; after reaching steady-state, MTY fluorescence is dictated only by the water-bath temperature; phase V, respiration stalled (anaerobiosis); temperature of the cell suspension medium (green trace) shifted by stepwise adjustments to water-bath temperature, followed by return to 38 °C. Measurements were carried out in a quartz chamber closed (capped cell) except for a 0.6 mm addition hole in the hand-made cap. The MTY fluorescence reached at the end of phase I was significantly different (n=10; ***) from the starting value of 50%, whilst the final value in phase IV was not. D, a, Linear increase of fluorescence of HEK293 cells (preloaded 10 min before trypsinization with 100 nM MTY) according to cell number (using cell protein concentration as surrogate parameter); b, Maximal rate of decrease of MTY fluorescence (%, blue circles; corresponding with mitochondrial warming) is not significantly affected by cell number, whereas initial fluorescence increase in presence of cyanide (%, green circles, corresponding with initial rate of mitochondrial cooling) is modulated by cell number (values at the three cell concentrations tested were significantly different from each other). E: a, HEK293 cells were made severely deficient for cytochrome *c* oxidase by culture (10 days) in the presence of ethidium bromide (EtBr; 1 µg/ml). Cytochrome *c* oxidase activity (blue circles) declined to a few percent of the activity measured at t=0, whilst citrate synthase activity (green circles) was little changed; b, The fluorescence of EtBr-treated HEK293 cells (10 days of EtBr treatment) pre-loaded with MTY (blue continuous line) does not decrease following suspension in oxygenated medium, whilst that of control HEK293 cells (blue dotted lines) follows the profile documented in Fig. 1C; in contrast to control cells (red dotted line), EtBr-treated HEK293 cells also do not consume appreciable amounts of oxygen (red continuous line).

We initially calibrated the response of MTY to temperature in solution. Its fluorescence at 562 nm (essentially unchanged by the pH of the solution buffer in the range 7.2 to 9.5; Fig. S2), decreased in a reversible and nearly linear fashion: a temperature rise from 34 to 60 °C decreased fluorescence by about 50%, whilst 82% of the response to a 3 °C shift at 38 °C was preserved at 50 °C (Fig. 1B, a, b). Using a thermostated, magnetically stirred, closable 750 µl quartz-cuvette fitted with an oxygen sensitive optode device [2] we simultaneously studied oxygen consumption (or tension) and changes in MTY fluorescence (Fig. 1C). Adherent cells were loaded for 20 min with 100 nM MTY, harvested and washed, then kept as a concentrated pellet at 38 °C for 10 min, reaching anaerobiosis in < 1 min. When cells were added to the oxygen-rich medium they immediately started to consume oxygen (red trace; Fig. 1C), accompanied by a progressive decrease of MTY fluorescence (blue trace; phase I; Fig. 1C). In the absence of any inhibitor, the fluorescence gradually reached a stable minimum (phase II). At that point, either due to a high temperature differential between the mitochondria and the surrounding cytosol (∼ 10°C) and/or changes in membrane permeability leading to decreased thermal insulation, the temperature of the probe-concentrating compartment appeared to reach equilibrium. We computed the energy released as heat by the respiratory chain in this experiment as 1.05-1.35 mcal/min, based on the measured rate of oxygen consumption (11.3 ± 1.8 nmol/min/mg prot), and assuming that heat accounts for the difference between the 52.6 kcal/mol released by the full oxidation of NADH and the 21 kcal/mol conserved as ATP under a condition of maximal ATP synthesis of 3 molecules per molecule of NADH oxidized. This should be sufficient to ensure the observed thermal equilibrium (∼50°C after 20 min) (See Appendix in Supplemental data). Once all the oxygen in the cuvette was exhausted (red trace) the directional shift of MTY fluorescence reversed (phase III), returning gradually almost to the starting value (phase IV). To calibrate the fluorescence signal *in vivo*, the temperature of the extra-cellular medium was increased stepwise (green trace). MTY fluorescence returned to the prior value when the medium was cooled again to 38 °C (Fig. 1C, phase V). This *in vivo* calibration was consistent with the response of MTY fluorescence in solution up to 44 °C, although further direct calibration steps *in vivo* are not possible without compromising cell viability. However, we confirmed that the response of MTY fluorescence to increased temperature deviates slightly from linearity *in vivo* in the same manner as in aqueous solution, namely that at maximal mitochondrial warming, extrapolated as being close to 50°C, the response to a 2 °C temperature shift is approximately 80% of that at 38 °C (Fig. S6). We therefore estimate the rise in mitochondrial temperature due to full activation of respiration as ∼10 °C (n=10, range 7-12 °C).

At the lowest (phase II) and highest fluorescence values (38°C, imposed by the water bath; phase IV), the signal was proportional to the amount of added cells, in a given experiment (Fig. 1D, a). Cell number did not affect the maximal rate of fluorescence decrease (computed from phase I). However, once anaerobic conditions had been reached, the initial rate of fluorescence increase (phase III, initial) was inversely related to the number of cells (Fig. 1D, b). To confirm that the observed fluorescence changes were due to mitochondrial respiration and not some other cellular process, we depleted HEK293 cells of their mtDNA with ethidium bromide (EtBr) to a point where cytochrome *c* oxidase activity was less than 2% of that in control cells (Fig. 1E, a, S1C). In EtBr-treated cells, no MTY fluorescence changes were observed under aerobiosis and cyanide treatment had no effect (Fig. 1E, b). Because MTY is derived from the membrane potential-sensitive dye rhodamine, whose fluorescence is essentially unaffected by temperature (Figure 1B, b), we investigated whether the observed changes in MTY fluorescence could be influenced by altered membrane potential, or by associated parameters. We took advantage of the fact that cyanide or oligomycin exert opposite effects on membrane potential (Fig. 2C, S1D) and compared the response of MTY fluorescence to these inhibitors (Fig. 2A,B). To avoid the possibly confounding effect of anaerobiosis, the quartz cuvette was kept uncapped in this experiment, with the oxygen tension rather than the rates of oxygen uptake being recorded (red traces). Once MTY fluorescence was stabilized (maximal mitochondrial heating), and the medium re-oxygenated, cyanide was added, causing a progressive increase in MTY fluorescence to the starting value (Fig. 2A). Note that, when cyanide was initially present, fluorescence changes and oxygen uptake were both abolished (Fig. 2A, dotted lines). Adding oligomycin in lieu of cyanide also decreased oxygen consumption and, as observed with cyanide, brought about a similar increase in MTY fluorescence (Fig. 2B). If added first, oligomycin progressively decreased oxygen uptake, abolishing the decrease in MTY fluorescence in parallel (Fig. 2B, dotted lines). Taken together, these experiments imply that electron flow through the respiratory chain (RC) rather than membrane potential or any related factor controls mitochondrial temperature. This conclusion is supported by examining the respective kinetics of membrane potential changes (tens of seconds) and MTY fluorescence changes (tens of minutes).

**Figure 2.**
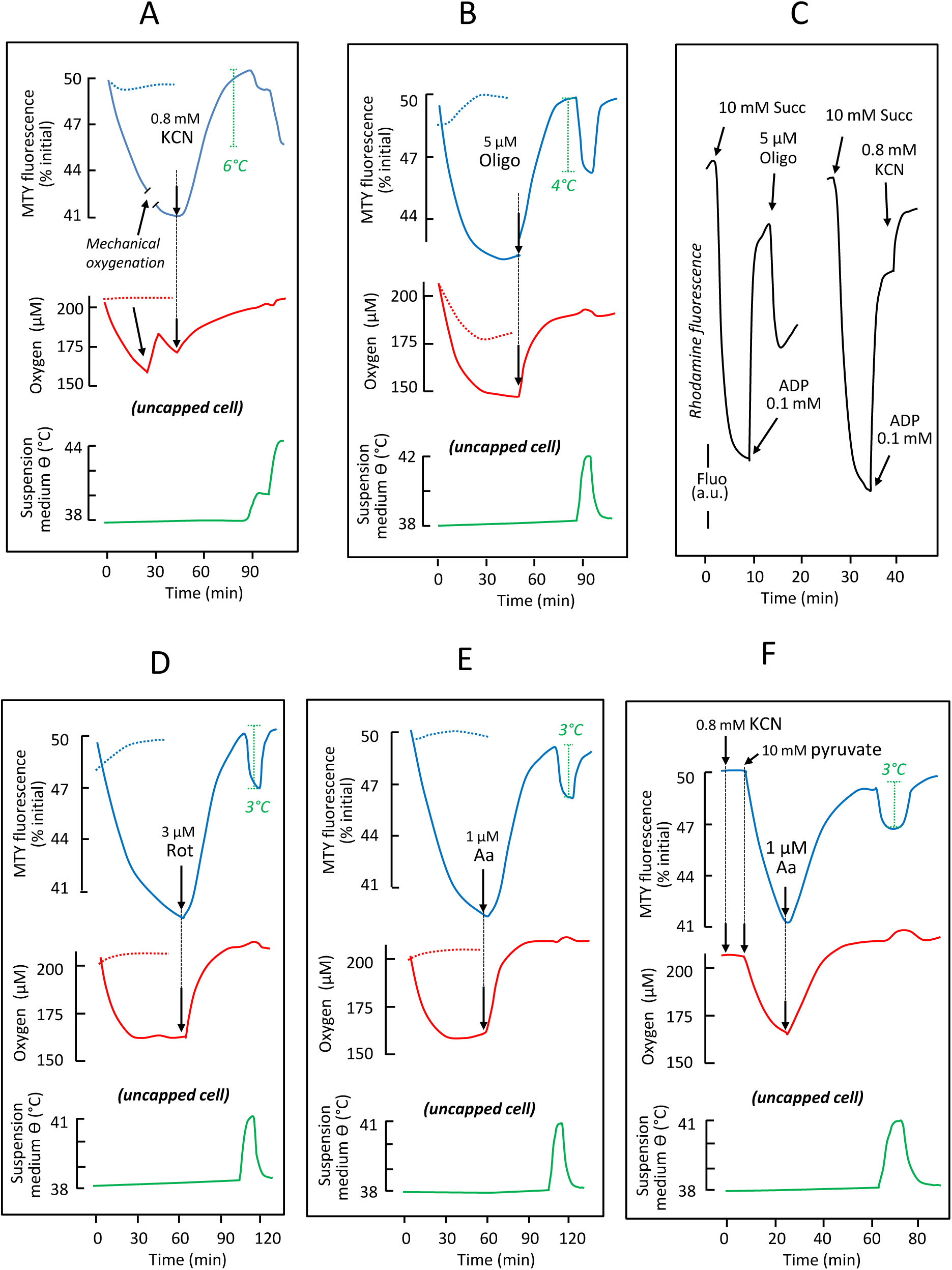
The rate of respiratory electron flow determines the temperature of mitochondria in intact HEK293 cells. A: The effect of 1 mM cyanide on MTY fluorescence (blue lines) and cell respiration (red lines), when added under aerobic conditions (continuous lines), or when present from the start of the experiment (dotted lines). Changes in the temperature of the cell suspension medium (green line), imposed by water bath adjustment, were used to calibrate the MTY fluorescence changes. B: The effect of 12.5 µM oligomycin on MTY fluorescence (blue lines) and oxygen tension (affected by cell respiration balanced by medium stirring) in the uncapped quartz-cuvette (red lines), when added to freely respiring cells (continuous lines), or when present from the start of the experiment (dotted lines). C: The effects of different inhibitors on rhodamine fluorescence, in digitonin (0.001%)-permeabilized HEK293 cells supplied with 10 mM succinate and 0.1 mM ADP as indicated. Under state 3 conditions 12.5 µM oligomycin and 0.8 mM KCN have qualitatively opposite effects on rhodamine fluorescence, used as an indicator of membrane potential (Δψ). The effects of 3 µM rotenone (D) and 1 µM antimycin (E) on MTY fluorescence and oxygen tension, plotted as for oligomycin in (B). F: The effect of adding pyruvate on MTY fluorescence (blue line) and oxygen uptake (red line) by KCN-inhibited HEK293cells. Temperature calibration (green line) of MTY fluorescence as in A. Note that in all experiments in which MTY fluorescence was measured, the value reached at the end of phase I was in all cases significantly different (n≥5; ***) from the starting value, whilst that in phase IV was not.

A quite similar effect was observed with two other respiratory inhibitors (Fig. S1D), affecting this time either RC complex I (CI; rotenone, Fig. 2D, S1D) or III (CIII; antimycin, Fig. 2E, S1D). Despite their different effects on the redox state of the various RC electron carriers, these inhibitors blocked oxygen uptake and again triggered an increase in MTY fluorescence. Importantly, these two inhibitors (and oligomycin) have been shown to trigger increased production of superoxide by the respiratory chain [3], but their effects on MTY fluorescence are similarly determined by oxygen consumption as for cyanide inhibition, which decreases superoxide production at complex III. This rules out any interference from superoxide in the observed MTY fluorescence changes. Taking advantage of cyanide removal from cytochrome *c* oxidase to form cyanohydrin in the presence of α-ketoacids under aerobiosis [4], we confirmed that the blockade of the respiratory chain did not result in MTY leakage from the mitochondria since pyruvate addition resulted in oxygen uptake resuming and MTY fluorescence decrease, both being inhibited by a further addition of antimycin (Fig. 2F). Leakage of the probe from mitochondria of other cell lines was reflected in a decreased ability of cyanide to restore MTY fluorescence to its initial value (Fig. S4C).

Note that changes in MTY fluorescence cannot be attributed to altered mitochondrial morphology, since 45 min of MTY treatment had no detectable effect on the mitochondrial network (Fig. S3i,j), nor did any of the inhibitors used in the above experiments (Fig. S3a-f). Importantly, the fluorescence of an endoplasmic reticulum (ER)-targeted version of the probe in HEK293cells and skin fibroblasts was essentially unaffected by the activity of the mitochondria when modulated by cyanide, pyruvate or antimycin (Fig. S7).

We next studied MTY probe behavior in HEK293 cells in which CI was inhibited by the addition of varying amounts of rotenone (Fig. 3A). The rate of change of MTY fluorescence was proportional to the residual respiratory electron flux whilst the maximal temperature, as judged by MTY fluorescence at equilibrium (phase II) was essentially unchanged (Fig. 3A; inset). We next tested the effect of expressing the cyanide-insensitive non-proton motive alternative oxidase from *Ciona intestinalis* (AOX; Fig. 3B, S1E), whose activity is unmasked in the presence of cyanide [5]. Before cyanide addition, the decrease in MTY fluorescence in AOX-expressing cells was similar to control cells, consistent with previous inferences that the enzyme does not significantly participate in uninhibited cell respiration [6]. However upon cyanide addition, AOX-endowed cells maintained low MTY fluorescence (Fig. 3B, blue trace), despite oxygen consumption decreasing by more than 50% (red trace). The increased ratio of heat generated to respiration is consistent with the predicted thermogenic properties of AOX. Subsequent addition of 0.1 mM propylgallate, which inhibits AOX, almost completely abolished the residual respiration and brought MTY fluorescence back to its starting value (corresponding to 38°C).

**Figure 3.**
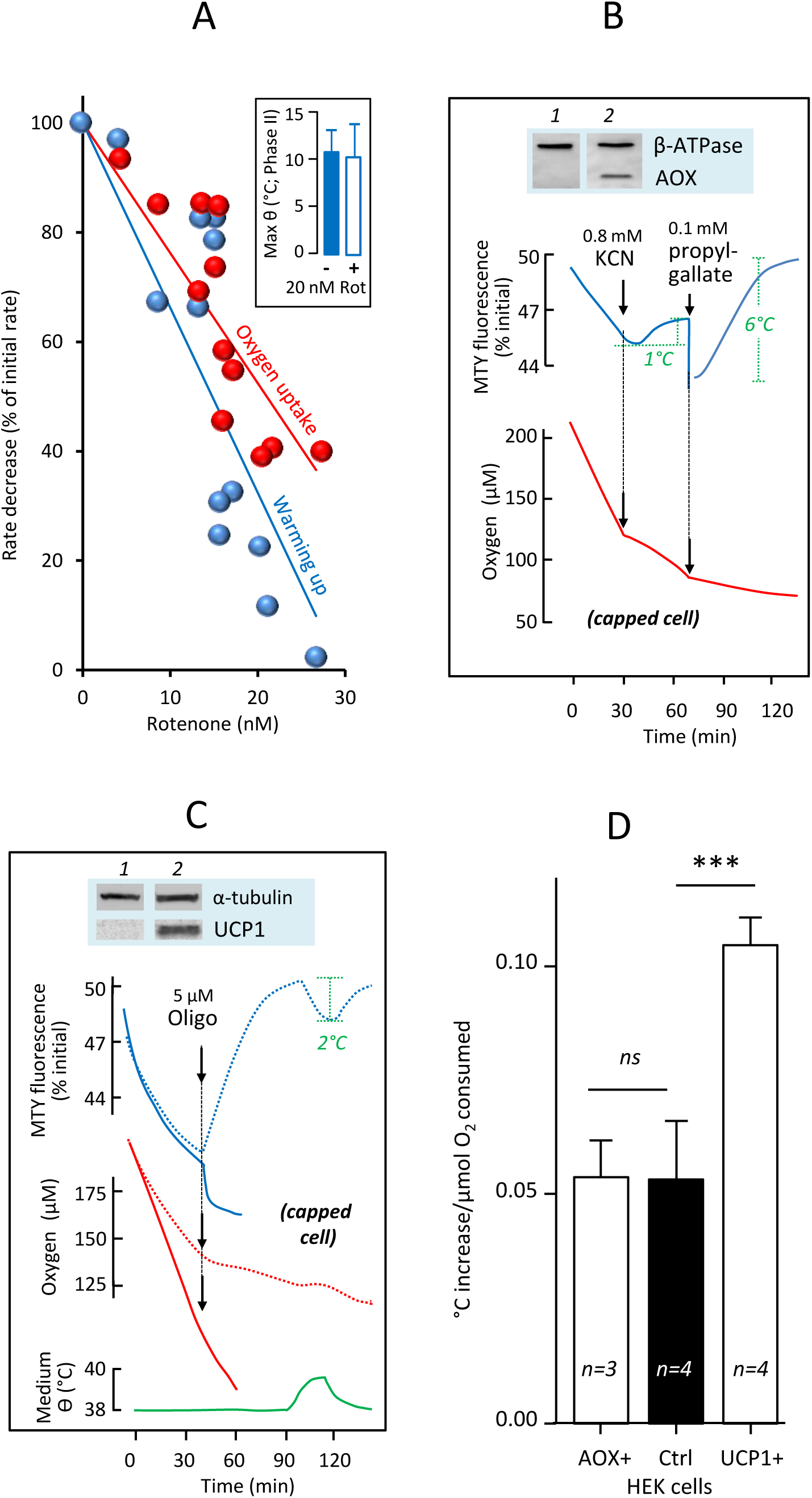
Effects on mitochondrial temperature of respiratory inhibitors, uncouplers and expression of heterologous mitochondrial proteins. A: Effect of variable rotenone addition to control HEK293 cells on the rates of oxygen uptake and fluorescence decrease of MTY. Rotenone was added at t=4 min; rates calculated from 4 to 7 min and expressed as a percent of initial rate. Inset: Maximal warming of HEK293 cell mitochondria in the absence or presence of 20 µM rotenone. B, C: Changes in MTY fluorescence (blue lines), cell respiration (B, C) (red lines) and temperature of cell suspension medium (green line), with additions of inhibitors as shown (KCN, n-propyl gallate) and/or oligomycin), alongside Western blots confirming expression or knockdown of the indicated genes: *C. intestinalis* alternate oxidase, AOX (B), UCP1 (C), alongside loading controls as indicated. Traces for control cells are shown by dotted lines. Note that in all experiments in which MTY fluorescence was measured, the value reached at the end of phase I was in all cases significantly different from the starting value, whilst that in phase IV was not (n=4; ***). (D), Computed from experiments using HEK293 cells endowed with AOX (AOX+), UCP1 (UCP1+) or control cells, initial increases of temperature (°C) per µmol oxygen consumed were compared and statistically analyzed by a one way ANOVA and Bonferroni’s multiple comparison test (n=3-4; means ± SD).

So as to circumvent the fact that we were not able to use chemical uncouplers with this probe [1], we used HEK293 cells engineered to express the uncoupling protein 1 (UCP1; Fig. 3C, S1F). As expected, UCP1 conferred an increased rate of respiration, which was only partially inhibited by oligomycin (red trace), and accompanied by an even greater drop in MTY fluorescence, equivalent to a temperature of about 12 °C above the cellular environment. HEK293 cells endowed with UCP1 also exhibited a faster rate of decrease of MTY fluorescence compared with control HEK293 cells, about two fold during the first 5 min (Fig. 3D).

The surprisingly high inferred mitochondrial temperatures prompted us to check the dependence on assay medium temperature of RC enzyme activities measured under V_max_ conditions in crude extracts, where mitochondrial membrane integrity is maintained (Fig. 4A, B). Antimycin-sensitive CIII, malonate-sensitive succinate cytochrome *c* reductase (CII+CIII) and cyanide-sensitive cytochrome *c* oxidase (CIV) activities all showed temperature optima at or slightly above 50 °C, whilst these activities tended gradually to decrease as the temperatures were raised further (Fig. 4A). This was not so for those enzymes whose activities can be measured *in vitro* only after osmotic disruption of both outer and inner mitochondrial membranes (Fig. 4B). Oligomycin-sensitive ATPase (CV) activity was optimal around 46 °C, whereas rotenone-sensitive NADH quinone-reductase (CI) activity declined sharply at temperatures above 38 °C. Interestingly after treatment at temperatures above 46 °C and 42 °C under conditions used for CI assay, the activities of CIV and CII of mitochondria were impaired as well, as revealed by native electrophoresis and in-gel activity (Fig. 4C).This strongly suggests a vital role for the inner mitochondrial membrane structure in the stabilization of the RC complexes at high temperature. We next analyzed the temperature profile of RC activity of primary skin fibroblasts. For CII+CIII, CIII and CIV (Fig. 4D), as well as CV (Fig. 4E) similar temperature optima were observed as in HEK293 cells, whilst MTY fluorescence (Fig. 4F) also indicated mitochondria being maintained at least 6-10 °C above environmental temperature.

**Figure 4.**
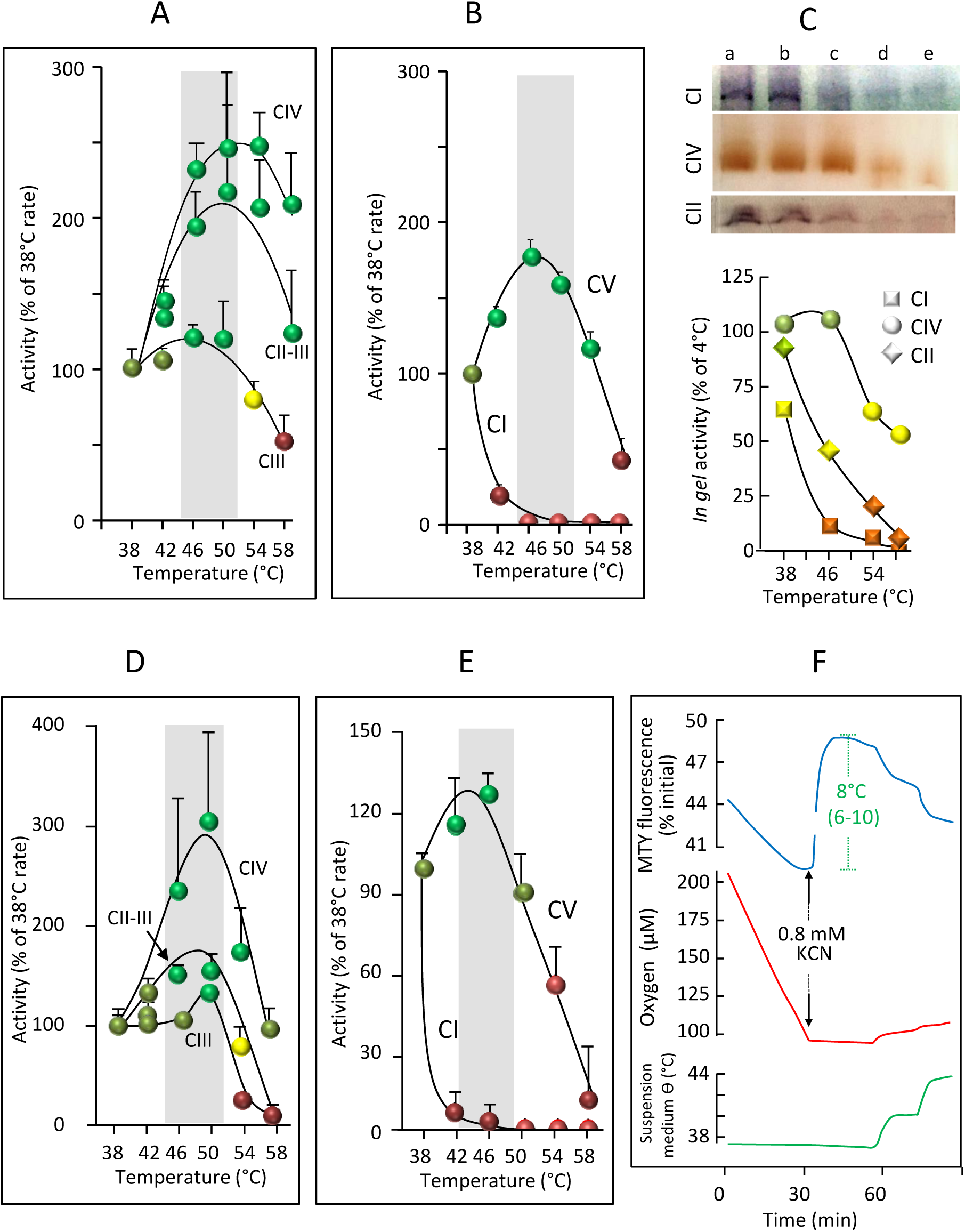
Effects of assay medium temperature on RC activities. A, D: Temperature profile of cytochrome *c* oxidase (CIV), malonate-sensitive succinate:cytochrome *c* reductase (CII+CIII) and antimycin-sensitive decylubiquinol:cytochrome *c* reductase (CIII) activity in (A) HEK293 cells, and (D) primary skin fibroblasts, after two freeze-thaw cycles, and (B, E) oligomycin-sensitive ATPase (CV) and rotenone-sensitive NADH:decylubiquinone reductase (CI) activity, after disruption of inner mitochondrial membrane in frozen (B) HEK293 cells and (E) primary skin fibroblasts, by osmolysis with water [27]. Colours denote optimal (green), minimal (red) and degrees of intermediate (pale greens, yellow) activity. Grey bars indicate optimal temperature range. C: CNE in-gel activities of CI, CIV and CII extracted from mitochondria previously incubated for 10 min at (a) 4 °C, (b) 37°C, (c) 42 °C, (d) 46 °C and (e) 55 °C, also plotted graphically (lower panel). F: Changes in MTY fluorescence (blue), cell respiration (red) and temperature of cell suspension medium (green), for primary skin fibroblasts, as denoted in Fig. 2A, with addition of KCN as shown.

## Discussion

Our findings raise numerous questions concerning the biochemistry, physiology and pathology of mitochondria. The physical, chemical and electrical properties of the inner mitochondrial membrane and of mitochondria in general, will need to be re-evaluated, given that almost all previous literature reflects experiments conducted far from our inferred physiological temperature. Traditional views of the lipid component of the respiratory membrane as a lake in which the RC complexes are floating resulting in a random-diffusion model of electron transfer or, more recently, as a sealant occupying the space between tightly packed proteins [7], need to be revised in favour of one that considers it more as a glue that maintains the integrity of the respiratory complexes.

A few years ago an intense debate took place on the actual possibility of maintaining temperature gradients in isolated cells considering the quite tiny volumes involved [8,9,10,11,12]. Largely based on theoretical consideration, it was suggested that additional factors must account for the large changes observed using thermosensitive-fluorescent probes [11]. For mitochondria, these potential factors include membrane potential changes (and related changes in pH, ionic gradients and matrix morphology), altered mitochondrial superoxide production, changes in probe conformation (especially for fluorescent protein probes) or probe leakage from mitochondria. Our study however suggests that none of these factors significantly influences MTY fluorescence under our experimental conditions.

On the other hand, the many unknowns regarding micro- and nano-scale physical parameters [12], render purely theoretical considerations questionable, in particular when considering the complex and dynamic structure of mitochondria. Most models have assumed mitochondria to be tiny, undifferentiated spheres, floating in an aqueous medium, the cytosol. This oversimplification would preclude significant temperature differences between mitochondria and the cytosol. However, mitochondria *in vivo* typically form a filamentous network, with considerable internal structure. Assuming MTY to be localized to the inner face of the inner membrane, or the adjacent pockets of matrix, the heated compartments would be juxtaposed to each other rather than to the colder cytosol. Moreover, compaction of the cytosol in domains rich in mitochondria, as observed in HEK293 cells, would also limit heat conduction out into the rest of the cell.

Lastly, the molecular heterogeneity of the various sub-mitochondrial compartments must be considered. The inner membrane is very rich in proteins (protein-to-lipid ratio 80:20, compared to 50:50 for the outer membrane), including those that are sources of heat, and has a distinct lipid composition including cardiolipins. The intermembrane space and the phospholipid-rich outer membrane might provide additional sources of insulation, with many relevant parameters unknown, including heat-conductance and geometry of the various compartments. Similar considerations may apply to hyperthermophile bacteria. These, in addition to thermal resistance of numerous compounds [13], have complex envelopes which have to act as insulators from extreme external environment as to maintain life-dependent membrane electrochemical gradients [14].

A 6 to 9 °C temperature shift between mitochondria and the surrounding cytosol, induced by the addition of an uncoupler (10 µM carbonyl cyanide-4-(trifluoromethoxy) phenylhydrazone), was recently reported in HeLa cells [15], using a genetically encoded, GFP-derived ratiometric fluorescent probe. Although carried under quite different conditions (confocal microscopy of a single cell), and without determining mitochondrial activity under these conditions, the data are consistent with our own findings.

Following our observations, the effects of respiratory dysfunction need to be reconsidered, to include those attributable to temperature changes, such as effects on membrane fluidity, electrical conductance and transport. RC organization into supercomplexes [16,17] should be re-examined at more realistic temperatures using methods other than CNE. Finally, whilst the subcellular distribution of mitochondria (e.g. perinuclear, or synaptic) has previously been considered to reflect ATP demand, mitochondria should also be considered as a source of heat, potentially relevant in specific cellular or physiological contexts, not just in specifically thermogenic tissues like brown fat. Furthermore, temperature differences should be considered as an additional possible dimension to the intracellular functional heterogeneity of mitochondria.

## Materials and Methods

### Cell culture

Human cells derived from Embryonic Kidney (HEK293), Hepatoma Tissue Culture (HTC-116) and from large cell lung cancer (NCI-H460) cells (American Type Culture Collection, Manassas, VA 20108 USA) were cultured in DMEM medium containing 5 g/l glucose supplemented by 2 mM glutamine (as Glutamax^™^), 10% foetal calf serum, 1 mM pyruvate, 100 µg/ml penicillin/streptomycin each. AOX-[18] or UCP-endowed [19,20]. The Trypan blue exclusion test was used to determine the number of viable and dead HEK cells [21]. Primary skin fibroblasts were derived from healthy individuals and grown under standard condition in DMEM glucose (4.5 g/l), 6 mM glutamine, 10% FCS, 200 µM uridine, penicillin/streptomycin (100 U/ml) plus 10 mM pyruvate.

### Immunoblot analyses and *in gel* enzyme activity assays

For the Western blot analysis, mitochondrial proteins (50 μg) were separated by SDS–PAGE on a 12% polyacrylamide gel, transferred to a nitrocellulose membrane and probed overnight at 4°C with antibodies against the protein of interest, AOX 1:10,000 [22], UCP1 1:10,000 [23]. Membranes were then washed in TBST and incubated with mouse or rabbit peroxidase-conjugated secondary antibodies for 2 h at room temperature. The antibody complexes were visualized with the Western Lightning Ultra Chemiluminescent substrate kit (Perkin Elmer). For the analysis of respiratory chain complexes, mitochondrial proteins (100 μg) were extracted with 6% digitonin and separated by hrCN-PAGE, on a 3.5–12% polyacrylamide gel. Gels were stained by in gel activity assay (IGA) detecting CI, CII and CIV activity as described [24].

### Staining procedures and life cell imaging

Cells (HEK293, MTC, NIC, primary skin fibroblasts) were seeded on glass coverslips and grown inside wells of a12 well-plate dish for 48 h in standard growth media at 37 °C, 5% CO2. The culture medium was replaced with pre-warmed medium containing fluorescent dyes, 100 nmol MitoTracker Green (Invitrogen M7514) and 100 nmol MitoThermo Yellow [1] or 100 nmol ER Thermo Yellow [25]. After 10 min the staining medium was replaced with fresh pre-warmed medium or PBS buffer and cells were observed immediately by Leica TCS SP8 confocal laser microscopy.

### Assay of mitochondrial respiratory chain activity

The measurement of RC activities was carried out using a Cary 50 spectrophotometer (Varian Australia, Victoria, Australia), as described [26]. Protein was estimated using the Bradford assay.

### Simultaneous spectrofluorometric, temperature, oxygen uptake assay

Detached sub-confluent HEK (NCI, HTS) cells (25 cm^2^ flask) or trypsinized sub-confluent skin fibroblasts (75 cm^2^ flask) were treated for 20 min with 100 µM MTY in 10 ml DMEM, and spun down (1,500 *g* x 5 min). The pellet is resuspended in 1 ml PBS, cells being next spun down (1,500 *g* x 5 min) and kept as a concentrated pellet. After anaerobiosis (checked by inserting an optic fiber equipped with an oxygen-sensitive fluorescent terminal sensor (Optode device; FireSting O_2_, Bionef, Paris, France) was established (10 min incubation of the pellet at 38°C;), cells (1 mg prot) were added to 750 µl of 38°C-thermostated medium. The fluorescence (excitation 542 nm, emission 562 nm for MTY; excitation 559 nm, emission 581 nm for ERTY), the temperature of the medium in the cuvette and the respiration of the intact cell suspension were simultaneously measured in a magnetically-stirred, 38°C-thermostated 1 ml-quartz cell in 750 µl of PBS using the Xenius XC spectrofluorometer (SAFAS, Monaco). Oxygen uptake was measured with an optode device fitted to a handmade cap, ensuring either closing of the quartz-cell yet allowing micro-injections (hole with 0.6 mm diameter) or leaving the quartz-cell open to allow for constant oxygen replenishment. Alternatively, untreated HEK293 cells (250 µg protein) were added to 750 µl of medium consisting of 0.25 M sucrose, 15 mM KCl, 30 mM KH_2_PO_4_ (pH 7.4), 5 mM MgCl_2_, EGTA 1 mM, followed by addition of rhodamine to 100 nM and digitonin to 0.01 % w/v. The permeabilized cells were successively given a mitochondrial substrate (10 mM succinate) and ADP (0.1 mM) to ensure state 3 (phosphorylating) conditions, under which either 6.5 µM oligomycin or 1 mM cyanide was added.

### Statistics

Data are presented as mean ± SD statistical significance was calculated by standard unpaired one-way ANOVA with Bonferroni post-test correction; a *p*< 0.05 was considered statistically significant (GraphPad Prism).

## Acknowledgements

This work was supported by French (ANR FIFA2-12-BSV1-0010 and ANR MITOXDRUGS-DS0403 to DC, PB, MR and PR) and European (E-rare Genomit to DC, PB, MR and PR) institutions and patient’s associations to PB and PR Association d’Aide aux Jeunes Infirmes (AAJI), Association contre les Maladies Mitochondriales (AMMi), Association Française contre l’Ataxie de Friedreich (AFAF), and Ouvrir Les Yeux (OLY).

## References

1. Arai S, Suzuki M, Park SJ, Yoo JS, Wang L, et al. (2015) Mitochondria-targeted fluorescent thermometer monitors intracellular temperature gradient. Chem Commun (Camb) 51: 8044–8047.

2. Benit P, Chretien D, Porceddu M, Yanicostas C, Rak M, et al. (2017) An Effective, Versatile, and Inexpensive Device for Oxygen Uptake Measurement. J Clin Med 6.

3. Boveris A (1977) Mitochondrial production of superoxide radical and hydrogen peroxide. Adv Exp Med Biol 78: 67–82.

4. Green DE, Williamson S (1937) Pyruvic and oxaloacetic cyanohydrins. Biochem J 31: 617–618.

5. Bahr JT, Bonner WD, Jr. (1973) Cyanide-insensitive respiration. II. Control of the alternate pathway. J Biol Chem 248: 3446–3450.

6. El-Khoury R, Dufour E, Rak M, Ramanantsoa N, Grandchamp N, et al. (2013) Alternative oxidase expression in the mouse enables bypassing cytochrome c oxidase blockade and limits mitochondrial ROS overproduction. PLoS Genet 9: e1003182.

7. Lenaz G, Baracca A, Barbero G, Bergamini C, Dalmonte ME, et al. (2010) Mitochondrial respiratory chain super-complex I-III in physiology and pathology. Biochim Biophys Acta 1797: 633–640.

8. Baffou G, Rigneault H, Marguet D, Jullien L (2014) A critique of methods for temperature imaging in single cells. Nat Methods 11: 899–901.

9. Sakaguchi R, Kiyonaka S, Mori Y (2015) Fluorescent sensors reveal subcellular thermal changes. Curr Opin Biotechnol 31: 57–64.

10. Kiyonaka S, Sakaguchi R, Hamachi I, Morii T, Yoshizaki T, et al. (2015) Validating subcellular thermal changes revealed by fluorescent thermosensors. Nat Methods 12: 801–802.

11. Baffou G, Rigneault H, Marguet D, Jullien L (2015) Reply to: “Validating subcellular thermal changes revealed by fluorescent thermosensors” and “The 10(5) gap issue between calculation and measurement in single-cell thermometry”. Nat Methods 12: 803.

12. Suzuki M, Zeeb V, Arai S, Oyama K, Ishiwata S (2015) The 10(5) gap issue between calculation and measurement in single-cell thermometry. Nat Methods 12: 802–803.

13. Stetter KO (1999) Extremophiles and their adaptation to hot environments. FEBS Lett 452: 22–25.

14. Oger PM, Cario A (2013) Adaptation of the membrane in Archaea. Biophys Chem 183: 42–56.

15. Nakano M, Arai Y, Kotera I, Okabe K, Kamei Y, et al. (2017) Genetically encoded ratiometric fluorescent thermometer with wide range and rapid response. PLoS One 12: e0172344.

16. Schagger H (2002) Respiratory chain supercomplexes of mitochondria and bacteria. Biochim Biophys Acta 1555: 154–159.

17. Wu M, Gu J, Guo R, Huang Y, Yang M (2016) Structure of Mammalian Respiratory Supercomplex I1III2IV1. Cell 167: 1598–1609 e1510.

18. El-Khoury R, Kaulio E, Lassila KA, Crowther DC, Jacobs HT, et al. (2016) Expression of the alternative oxidase mitigates beta-amyloid production and toxicity in model systems. Free Radic Biol Med 96: 57–66.

19. Oelkrug R, Goetze N, Exner C, Lee Y, Ganjam GK, et al. (2013) Brown fat in a protoendothermic mammal fuels eutherian evolution. Nat Commun 4: 2140.

20. Keipert S, Jastroch M (2014) Brite/beige fat and UCP1 -is it thermogenesis? Biochim Biophys Acta 1837: 1075–1082.

21. Strober W (2001) Trypan blue exclusion test of cell viability. Curr Protoc Immunol Appendix 3: Appendix 3B.

22. Hakkaart GA, Dassa EP, Jacobs HT, Rustin P (2006) Allotopic expression of a mitochondrial alternative oxidase confers cyanide resistance to human cell respiration. EMBO Rep 7: 341–345.

23. Jastroch M, Hirschberg V, Klingenspor M (2012) Functional characterization of UCP1 in mammalian HEK293 cells excludes mitochondrial uncoupling artefacts and reveals no contribution to basal proton leak. Biochim Biophys Acta 1817: 1660–1670.

24. Wittig I, Karas M, Schagger H (2007) High resolution clear native electrophoresis for ingel functional assays and fluorescence studies of membrane protein complexes. Mol Cell Proteomics 6: 1215–1225.

25. Arai S, Lee SC, Zhai D, Suzuki M, Chang YT (2014) A molecular fluorescent probe for targeted visualization of temperature at the endoplasmic reticulum. Sci Rep 4: 6701.

26. Rustin P, Chretien D, Bourgeron T, Gerard B, Rotig A, et al. (1994) Biochemical and molecular investigations in respiratory chain deficiencies. Clin Chim Acta 228: 35–51.

27. Chretien D, Benit P, Chol M, Lebon S, Rotig A, et al. (2003) Assay of mitochondrial respiratory chain complex I in human lymphocytes and cultured skin fibroblasts. Biochem Biophys Res Commun 301: 222–224.

